# Green microwave-assisted extraction of *Euphorbia guyoniana:* Optimization, metabolite profile and *in vivo* anti-inflammatory potential

**DOI:** 10.1101/2024.05.10.593649

**Authors:** Halima Meriem Issaadi, Yacine Nait Bachir, Alaaeddine Ben Naama, Imad Aiter, Kornél Szőri, Attila Hunyadi

## Abstract

This study aimed to optimize microwave-assisted extraction of phenolic compounds from undervalued traditional plant *Euphorbia guyoniana* (Boiss. & Reut.) using central composite design of response surface methodology. The independent variables were extraction time (*x_1_*: 5 – 25 min), ethanol concentration in the extractive solvent (*x_2_*: 30 – 70%), microwave power (*x_3_*: 180 – 800 Watt) and feed-to-solvent ratio (*x_4_*: 1:7.5 – 1:17.5) while dependent variables were total phenolic content (TPC) and total flavonoid content (TFC). Extract obtained by using the optimal extraction parameters was evaluated for its *in vivo* anti-inflammatory activity by the carrageenan-induced paw edema model and was subjected to RP-HPLC-PDA-ESI-MS analysis to investigate the presence of phenolic compounds. The optimal conditions for highest TPC (377.22 ± 5.42 mg GAE/100g) and TFC (184.40 ± 1.18 mg QE/100g) were obtained at extraction time of 25 min, ethanol concentration of 40.57%, microwave power of 450 Watt and feed-to-solvent ratio of 1:17.5. The quadratic models significantly (p < 0.0001) fitted the experimental data with R^2^ values of 0.984 and 0.970 for TPC and TFC, respectively. Optimal extract of *Euphorbia guyoniana* significantly higher inhibited carrageenan induced inflammation with a concentration of 50 mg/kg (75,98%) when compared with reference anti-inflammatory drug ibuprofen (43,77%). Finally, above the previously reported phenolic constituents, i.e, quercetin and kaempferol derivatives, hydroxycinnamates have been identified for the first time in *Euphorbia guyoniana* plant extract.

## 1. Introduction

*Euphorbia guyoniana* (Boiss. & Reut.) is a perennial dark green herb with a creeping underground stem, bright green 30-100 cm in height from the genus *Euphorbia* of the Euphorbiaceae plant family [**1, 2**]. Over 1600 species belong to the genus *Euphorbia* and are native to tropical and subtropical regions of Asia, Africa, Central and South America [**2**]. *Euphorbia guyoniana* is an endemic plant to Algeria, Tunisia, Libya and Morocco and is located in sandy regions, pre-desert and in the north of Sahara. The plant is used in the ethno-medicine of North African countries: in Tunisia, the plant is widely used as an antitussive and analgesic remedy and in the Saharan region of Algeria and Morocco as a wart remover, and for the treatment of venomous scorpion bites [**3, 4**]. However, to the best of our knowledge, only a few scientific studies dealt with the medical importance of the aerial part of this plant and its bioactive constituents responsible for its folk use. The aerial part of *E. guyoniana* was reported to exert antimicrobial [**5, 6, 7, 8**], antifungal [**9**], cytotoxic [**7, 10**], antioxidant [**10, 11**] and antihyperglycemic [**12**] activities. Phytochemical studies on the aerial part of *E. guyoniana* revealed the presence of jatrophane diterpenes [**10, 13**], polyester diterpenes [**14**], flavonoids [**11, 15**] and an alkaloid [**16**].

In recent years, traditional methods of extracting medicinal plants have been increasingly replaced by novel unconventional methods, most of which have been claimed to be more effective in terms of yield, solvent usage, extraction time and efficiency [**17**]. Microwave-assisted extraction (MAE) has shown a notable advantage over other modern methods, including significantly reduced processing time and improved yields [**18**]. The efficiency of microwave-assisted extraction is influenced simultaneously by several factors such as microwave power, irradiation time, sample-to-solvent ratio, temperature, etc. [**19**]. To understand the effect of these factors and their possible interactive impact on extraction outcomes, response surface methodology (RSM) has been introduced and widely used nowadays [**20, 21**]. RSM is a powerful mathematical statistical technique that uses quantitative data from experimental design to show a correlation between extraction parameters and their responses at different optimum levels and to predict the optimal conditions for the extraction process.

In this study, response surface methodology using central composite design (CCD) in five different coded levels was employed for the experimental planning of the green microwave extraction of the aerial part of *Euphorbia guyoniana*. To the best of authors’ knowledge, there is no previous report in the literature on the MAE of *Euphorbia guyoniana* on the total phenolic content and total flavonoid content. Using the optimized extract of *E. Guyoniana*, an *in vivo* anti-inflammatory using the rat paw-edema assay study was conducted. Finally, the phenolic bioactive compounds were screened by Reverse-Phase High Performance Liquid Chromatography coupled to a Mass Spectroscopy detector (RP-HPLC-PDA-ESI-MS) in positive ionisation.

## 2. Results and discussion

RSM is a statistical approach that can model and optimize a process using fewer and more informative experiments than the classical “one-factor-at-a-time” approach or the use of full factorial designs, and it indicates the possible influences of some variables on others. In this study, experiments conducted according to the RSM-CCD included 27 runs. The experimental results for total phenolic content (TPC) and total flavonoid content (TFC) of *Euphorbia guyoniana* extracts obtained under different combinations of independent variables (extraction time, ethanol concentration, microwave power and feed-to-solvent ratio) are presented in Table 6. The values ranged from 191.89 mg GAE/100g (experiment 6) to 301.47 mg GAE/100g (experiment 3) and from 108.64 mg QE/100g (experiment 19) to 159.57 mg QE/100g (experiment 17) respectively, which implies that the selected and studied factors have a real influence on the quantity of these bioactive constituents in the extract.

### 2.1 Optimization of microwave-assisted extraction (MAE) and models fitting

The model fittings for TPC and TFC of *Euphorbia guyoniana* extracts are essential in evaluating how the response surface methodology mathematical models can predict the variance and show the correlation between the extraction parameters of MAE and chemical composition of the extracts. The analysis of variance of the CCD for the models fitting and the correlations between the predicted and experimental actual values are shown in Table 1 and (Fig 1), respectively.

**Fig 1.**
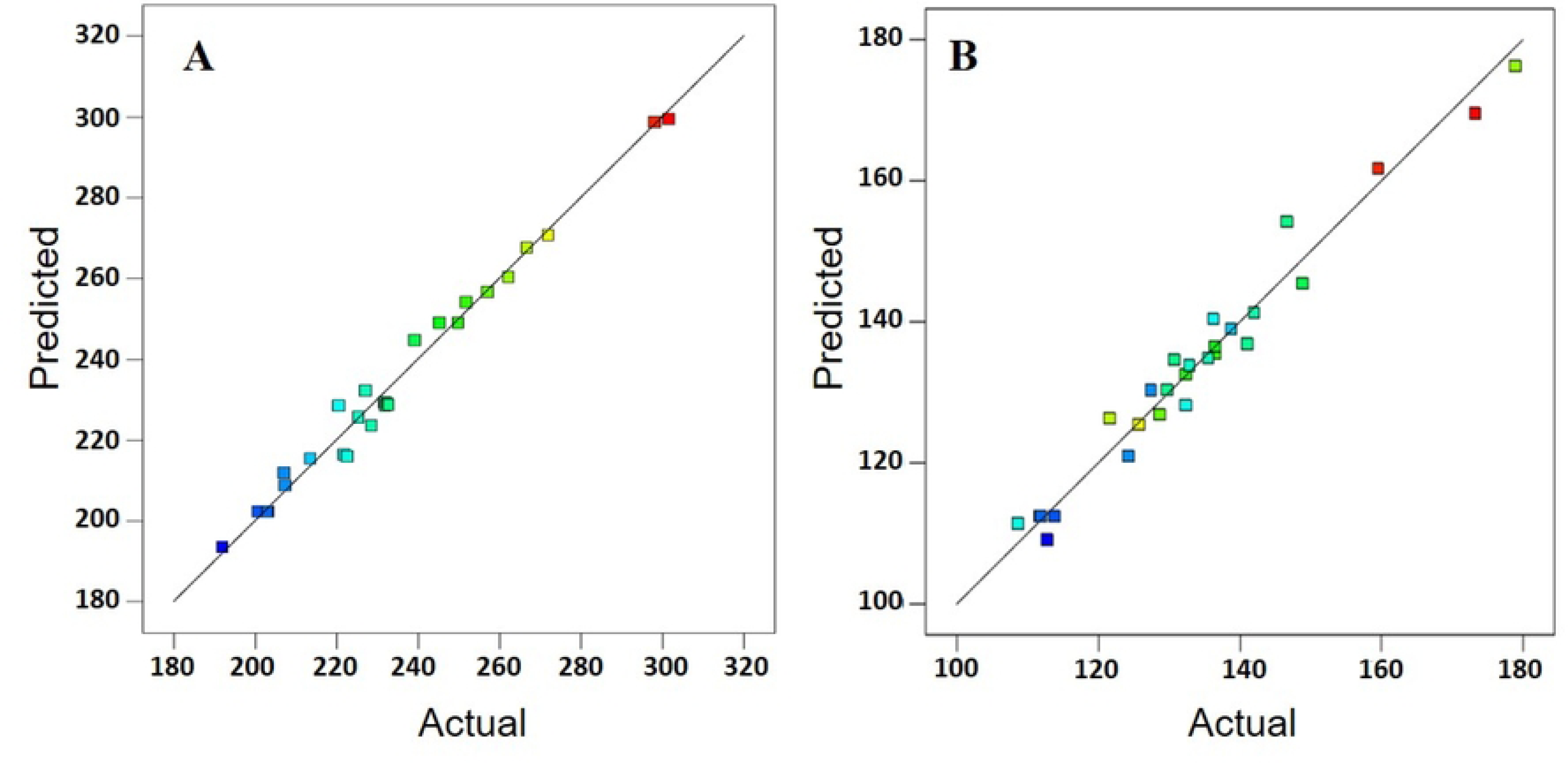
Correlation between predicted and experimental actual total phenolic content (A) and total flavonoid content (B)

**Table 1.**
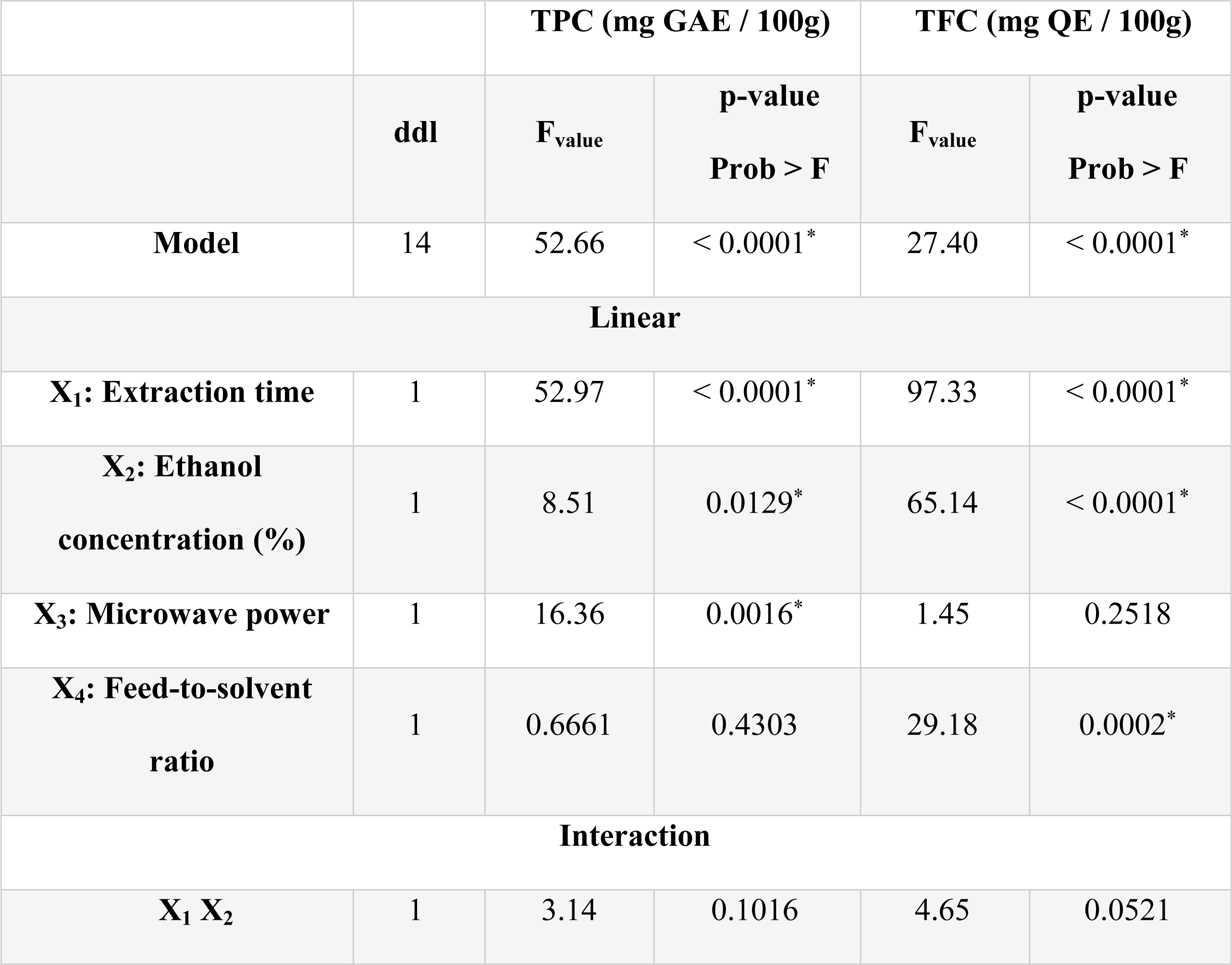
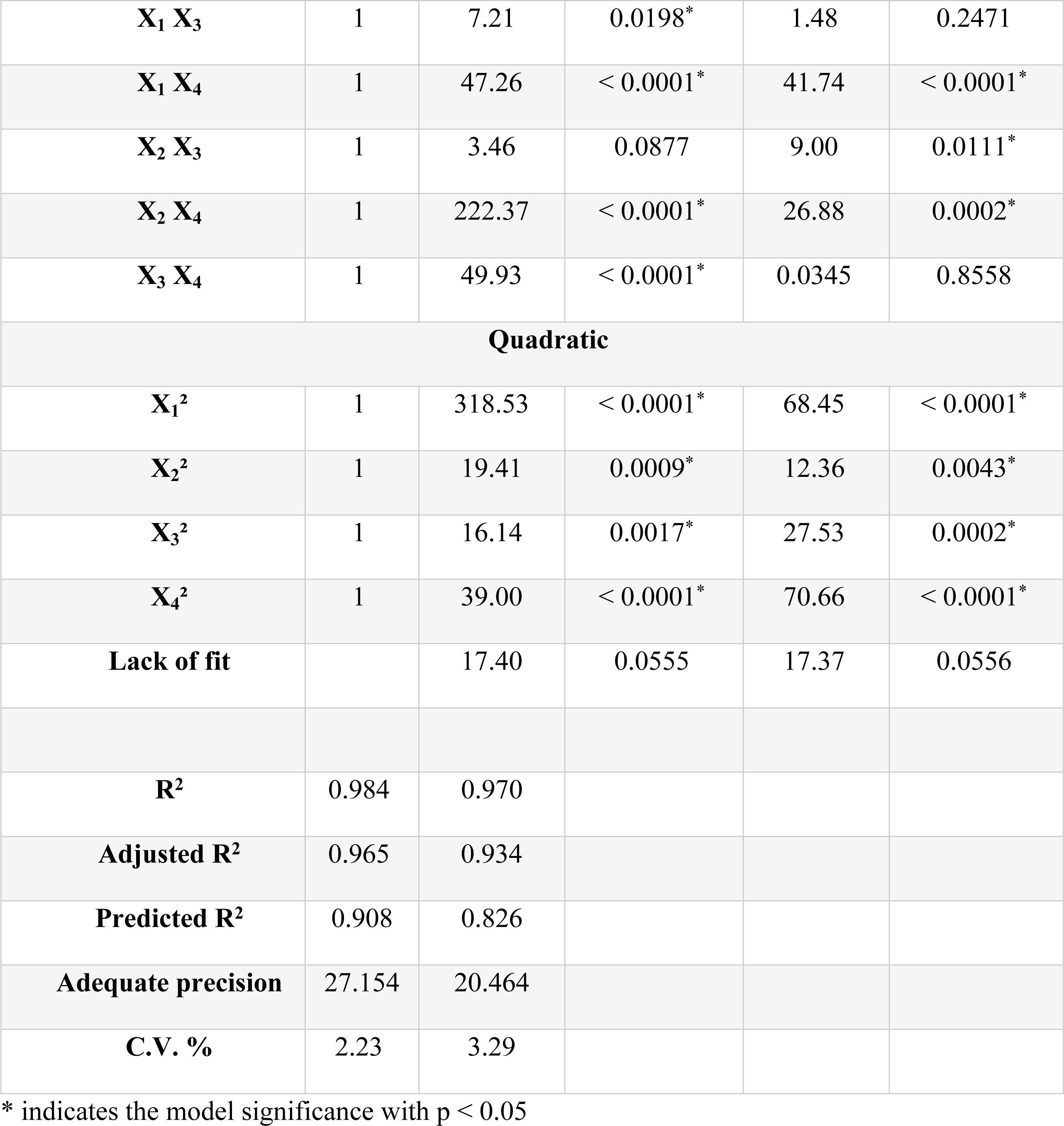
Experimental results analysis of variance.

Both analysis of variance and the statistical parameters shown in Table 1 indicate that quadratic second-order polynomial models were estimated as the most suitable for fitting all the experimentally obtained responses without any transformations and adequate to predict total phenolic and total flavonoid contents extracted by microwave procedure from *Euphorbia guyoniana*.

For the TPC the p-value of the model was < 0.0001, indicating that the model was significant and the p-value for the lack of the model was calculated to be 0.0555 showing the non-significance (p > 0.05) of the model lack of fit. Also, the coefficient of determination (R^2^) was 0.984, demonstrating a 98.40% match between the predicted and actual data. The predicted R^2^ of 0.908 was in reasonable agreement with the adjusted R^2^ of 0.965 and the adequate precision which measure signal to noise ratio was greater than 4. This indicates an adequate signal allowing the model to be used to navigate the design space. Finally, the coefficient of variation (C.V.) which evaluates the relative closeness of the predictions to the actual values was estimated to be 2.23%. This low value illustrates a high degree of precision of the experimental values which allows for the development of an appropriate model.

In the same way, for the model fitting for TFC, p-value of the model, p-value for lack of fit, R^2^, adjusted R^2^, predicted R^2^, adjusted R^2^, adequate precision, and C.V. were calculated to be < 0.0001, 0.0556, 0.970, 0.934, 0.826, 20.464 and 3.29% respectively confirming the significance and reliability of the model in predicting the TFC yield.

Lastly, a strong linear correlation was observed between the predicted and experimental total phenolic content (A) and total flavonoid content (B) as presented in (Fig 1). Uniform distribution of the experimental values around the fitted line, and the high R^2^ values obtained indicate the reasonable precision between the data and the model (All experimental and predicted values are presented in Supporting S1 Table).

### 2.2 The influence of extraction factors on TPC and TFC of *Euphorbia guyoniana* extracts

According to the results of ANOVA presented in Table 1 and considering the significant p-value of the studied parameters, TPC recovery was affected the most significantly by *x_1_* (extraction time, p < 0.0001). This was followed by *x_3_* (microwave power, p = 0.0016) and *x_2_* (concentration of ethanol in the extractive solvent, p = 0.0129). Interestingly, the linear feed-to-solvent ratio *x_4_* parameter did not affect (p > 0.05) TPC values; however, its interaction with all other parameters i.e. *x_1_x_4_*, *x_2_x_4_*, *x_3_x_4_* was highly significant (p < 0.0001). Finally, all the quadratic terms impacted positively the response, i.e., *x_1_*² and *x_4_*² (p < 0.0001), *x_2_*² (p = 0.0009) and *x_3_*² (p = 0. 0017).

The mathematical quadratic modeling of the extraction of TPC as a function of the different parameters is represented by Eq. (1), it shows the relation between the significant extraction variables and the TPC as response:

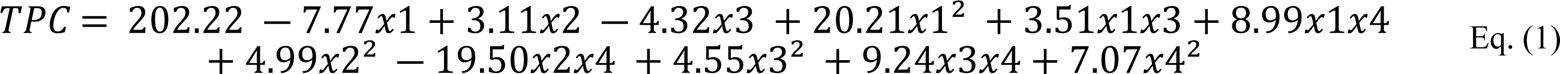

In Table 1, it can be clearly seen that TFC recovery was affected by three linear parameters *x_1_* (extraction time) *x_2_* (concentration of ethanol in the extractive solvent) and *x_4_* (feed-to-solvent ratio). Three interaction terms i.e. *x_1_x_4_*, *x_2_x_3_* and *x_2_x_4_* were significant (p < 0.05). As for the TPC recovery, all quadratic terms presented a significant effect on the response: *x_1_*² and *x_4_*² (p < 0.0001), *x_3_*² (p = 0.0002) and *x_4_*² (p = 0.0043).

The mathematical modeling of the extraction of TFC for the response variable was expressed by the following equation Eq. (2) in the form of coded values:

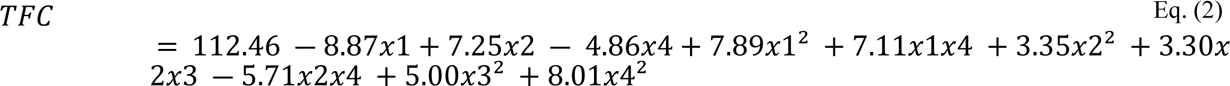

### 2.3 Experimental validation of the optimization

In order to optimize the obtained TPC and TFC, an optimization study in the larger variable domain from -α to +α was conducted. With the aim to not overstep limitations of the microwave, i.e., discontinuous microwave power values, it was decided to fix the microwave parameter to the central point 0. Further, to avoid the evaporation of extractive solvent (phenomenon observed when the lowest sample-to-solvent ratio was applied), the parameter *x_4_* (sample-to-solvent ratio) was adjusted to “maximize”. All other independent variables were adjusted to “in range” and to “maximize” for all responses simultaneously. Different possible combinations with desirability equal to 1 were proposed by the software, however they were not experimentally possible. A combination with a desirability of 0.951 was selected.

The values of extraction factors that maximized the recovery of TPC and TFC from aerial parts of *E. guyoniana* in the microwave reactor were 25 min of extraction time, 40.57% of ethanol in the solvent extraction, 450 Watt and 1:17.5 of sample-to-solvent ratio. The optimized conditions were experimentally validated by conducting three replicate extractions. Under these conditions the total phenolic content was estimated to be 376.16 mg GAE/100g while the measured value was 377.22 ± 5.42 mg GAE/100g. Similarly, the total flavonoid content was estimated to be 188.46 mg QE/100g while the measured value was 184.40 ± 1.18 mg QE/100g. The values of the TPC and TFC content obtained by experimental extractions were close to those predicted by the models (Table 2), demonstrating that the developed RSM models fit well for the extraction of bioactive compounds from *E. guyoniana* under optimal MAE condition and that they exhibit high predictive capacity and repeatability.

**Table 2.** The predicted and experimental values at the optimal settings for microwave-assisted extraction of total phenolics content (TPC) and total flavonoids content (TFC) from *Euphorbia guyoniana*.

| Optimized extraction conditions |  | TPC<br>(mg GAE / 100g) |  | TFC<br>(mg QE / 100g) |  |
| --- | --- | --- | --- | --- | --- |
|  |  | Actual | Predicted | Actual | Predicted |
|  |  | value | value | value | value |
| Extraction time (min) | 25 | 377.22 ±<br>5.42 | 376.16 | 184.40 ±<br>1.18 | 188.46 |
| Ethanol concentration (%) | 40.57 |  |  |  |  |
| Microwave power (Watt) | 450 |  |  |  |  |
| Feed-to-solvent ratio (g/ml) | 1:17.5 |  |  |  |  |

### 2.4 Analysis of response surface methodology

To visualize and facilitate understanding the mutual interactions between the studied independent parameters on TPC and TFC yields, the three-dimensional (3D) response surface plots are shown in (Figs 2 and 3), respectively. They are generated by plotting responses against two independent parameters while keeping other parameters at zero levels.

**Fig 2.**
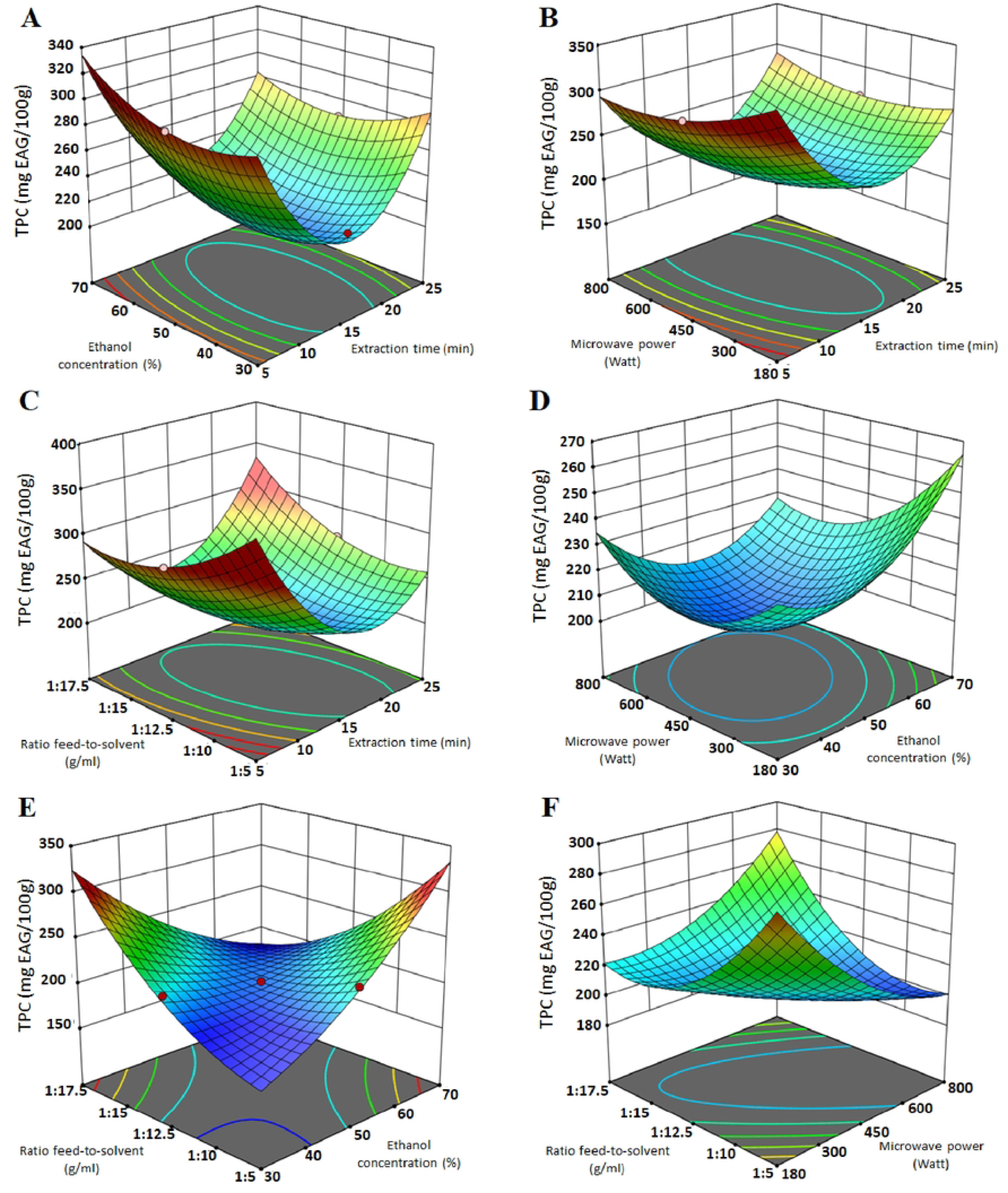
3D response surface plots for total phenolic content (TPC) of *Euphorbia guyoniana* extract in function of extraction time, concentration of ethanol, microwave power and feed-to-solvent ratio.

**Fig 3.**
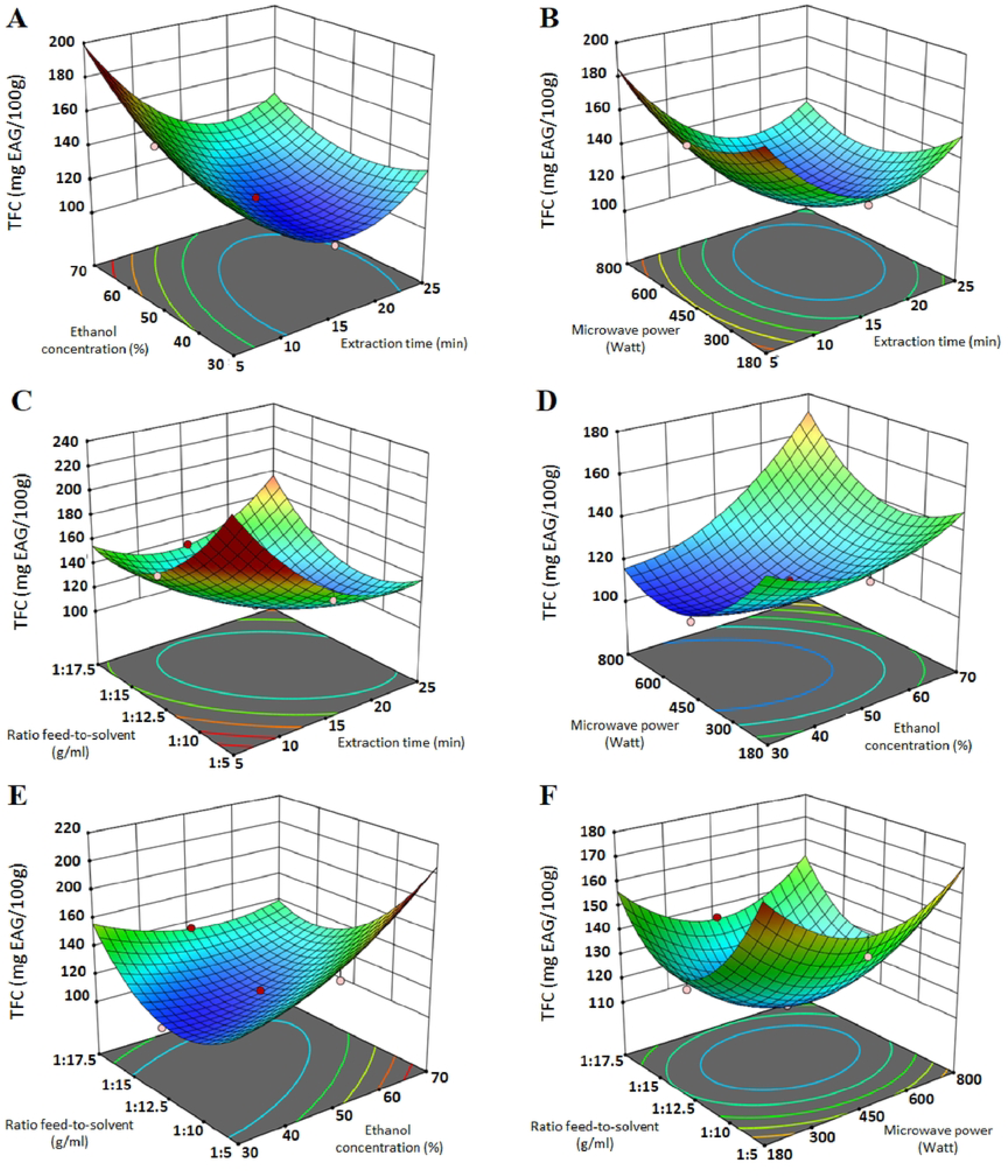
3D response surface plots for total phenolic content (TFC) of *Euphorbia guyoniana* extract in function of extraction time, concentration of ethanol, microwave power and feed-to-solvent ratio.

The interaction of the parameter feed-to-solvent ratio with all studied parameters presented a highly significant effect. Indeed, in all the cases, taking extreme values of the feed-to-solvent ratio had a positive impact on bioactive compounds levels (Figs 2 and 3 C, E, F). In this context, on one hand, the use of a low ratio increases the extraction efficiency of bioactive compounds, but may also lead to solvent evaporation. On the other hand, a high feed-to-solvent ratio increases the contact time between the solvent and the plant matrix, leading to a higher yield of bioactive compounds; however in the extraction process at an industrial scale, a balance needs to be found between minimizing the high costs of solvent consumption and its waste and maximizing extraction yield. (Figs 2 and 3 C) show that the decrease of feed-to-solvent ratio along with extraction time impacted positively the levels of TPC and TFC. This trend is maintained when feed-to-solvent ratio and extraction time are simultaneously increased. This observations suggest that the planned time extraction choice is therefore closely related to the feed-to-solvent ratio. In our case, the use of feed-to-solvent ratio of 17.5 g of *E. Guyoniana*/ml automatically led us to increase the extraction time to allow the temperature inside the extraction solvent-vegetable matrix mixture to increase and be evenly distributed.

Based on the statistical results, the significant negative linear effect and positive quadratic effect of extraction time, on both studied responses (TPC and TFC), suggest that the relationship between the independent variable and the dependent variables is not linear but rather follows a U-shaped curve. When the extraction time increases, the effect on TPC and TFC first decreases, reaches a peak, and then starts to increase. This phenomenon could be because extended irradiation time, especially under the influence of increased temperature in MAE, may cause enzymatic degradation or oxidation of polyphenols, resulting in a loss of observed antioxidant activity. It was already reported that highly numbered hydroxyl-substituted phenolic compounds (e.g., tannins) and those that are sensitive to elevated temperature (e.g., anthocyanins) may not be suitable to be extracted by MAE [**22**]. Accordingly, MAE has been found useful in the extraction of short-chain compounds like phenolic acids and flavonoids [**23**] which is in total accordance with the results of our study where mainly hydroxycinnamic acids and flavonol glycosides where identified from the studied plant.

The interaction of microwave power with the three applied parameters always impacted the yields of phenolic compounds and flavonoids in the same way, i.e. in all cases, the decrease in microwave power had a positive impact on the levels of bioactive compounds. Increasing the microwave power led to an increase in temperature, which may enhance the yield of phenolic compounds by improving their solubility due to increased solvent accessibility. However, applying excessively high temperatures has been found to degrade phenolic compounds, resulting in reduced levels [**18**]. Also, it was reported that the application of microwaves equal to or greater than 800 Watts can lead to the carbonization of plant material [**24**]. These conclusions are consistent with a study on microwave-assisted extraction conducted on *Euphorbia hirta*, where the optimal microwave power was determined to be 400 Watt [**25**]. Considering this, in the current study we decided to fix the microwave parameter to the central point 0 corresponding to 450 Watt for the optimization procedure.

Phenolic compounds are extracted from various plant materials using solvents like alcohols, acetone, diethyl ether and ethyl acetate, often mixed with different proportions of water. Ethanol is a good solvent for polyphenolic extraction as it is safe for human consumption as recommended by the Food and Drug Administration (FDA). Also, using ethanol with different proportions of water causes a decrease in the dielectric constant of the extraction medium, which enables easier separation of the solvent molecules [**26**]. Thereby, in this study, ethanol with different proportion of water has been used. The ethanol percentage had a significant positive effect on TPC and TFC indicating that with the increase in ethanol concentration, the bioactive contents would increase (Figs 2 and 3). It is already known that the highest yields of flavonoids are achieved when using aqueous-ethanol as the extraction medium based on the fact that ethanol forms non-covalent bonds with flavonoids, facilitating their diffusion into the solvent [**27**].

### 2.5 Inhibitory effect of the optimal *Euphorbia guyoniana* extract on carrageenan-induced paw edema rats

To evaluate the anti-inflammatory activity of the optimal *Euphorbia guyoniana* extract, the carrageenan-induced paw edema model was used. The model involved subcutaneously administering carrageenan to rats which caused an increase in paw size due to edema.

As shown in Table 3, sub-plantar injection of the carrageenan induced a significant time dependency increase in paw edema in the three groups. The increase in paw edema was calculated for each group as compared to the pre-injection values: Group 1 composed of 6 rats receiving no treatment (negative control), Group 2 consisting of 6 rats receiving the anti-inflammatory ibuprofen at a dose of 50 mg/kg (positive control) and Group 3 composed of 6 rats receiving 50 mg/kg of *Euphorbia guyoniana* extract. The percentages of inhibition of paw edema for the two treated groups i.e. Group 2 receiving the anti-inflammatory ibuprofen (positive control) and Group 3 treated with *Euphorbia guyoniana* extract were of 43.77% and 75.98%, respectively. Thus, rats treated with the extract showed a higher percentage of inhibition of paw edema than that obtained with a reference treatment. To the best of our knowledge, this is the first report on the anti-inflammatory potential of *E. guyoniana*. These conclusions corroborate the use of this plant in traditional Moroccan and Algerian medicine as a local anti-inflammatory for the treatment of venomous bites and stings [**3, 4**].

**Table 3.** Assessment of the anti-inflammatory activity of *E. guyoniana* in carrageenan induced paw edema.

|  | Treatment | Paw<br>volume<br>(0h) | Paw<br>volume<br>(1h) | Paw<br>volume<br>(2h) | Paw<br>volume<br>(3h) | Paw<br>volume<br>(4h) | %<br>Inhibition<br>of Edema |
| --- | --- | --- | --- | --- | --- | --- | --- |
| **Group 1** | Physiological<br>saline | 1,27 ±<br>0,04 | 1,74 ±<br>0,11 | 2,45 ±<br>0,26 | 3,00 ±<br>0,25 | 3,11 ±<br>0,26 | - |
| **Group 2** | Ibuprofen –<br>50 mg/kg<br>(positive<br>control) | 1,25 ±<br>0,03 | 1,43 ±<br>0,06 | 1,63 ±<br>0,10** | 1,85 ±<br>0,11** | 1,87 ±<br>0,11** | 43,77 |
| **Group 3** | *E. guyoniana* –<br>50 mg/kg | 1,26 ±<br>0,02 | 1,34 ±<br>0,03 | 1,41 ±<br>0,05* | 1,45 ±<br>0,04** | 1,47 ±<br>0,04** | 75,98 |

Each value is presented as mean ± SEM (n = 6 rats; one-way ANOVA followed by *Post hoc* Bonferroni test). \*: p < 0.05, \*\*: p < 0.001 compared with the negative control group

### 2.6 Phenolic compounds profile via RP-HPLC-PDA-ESI-MS

Phenolic compounds were analyzed by RP-HPLC-PDA-ESI-MS in the extract from the aerial part of *Euphorbia guyoniana* obtained at the optimized conditions of microwave assisted extraction. The extract was subjected to sequential fractionation using organic solvents of increasing polarity resulting in three less complex fractions. All fractions were analyzed successively and their characteristic chromatograms are presented in supporting information (S1 Fig, S2 Fig and S3 Fig). A total number of 12 phenolic compounds were tentatively identified by comparing their spectral data with those available in the literature. The identified phenolic compounds, their retention times, maximal UV absorbance, molecular ion and main fragment ions are presented in Table 4. The analysis also included ten unidentified compounds.

**Table 4.**
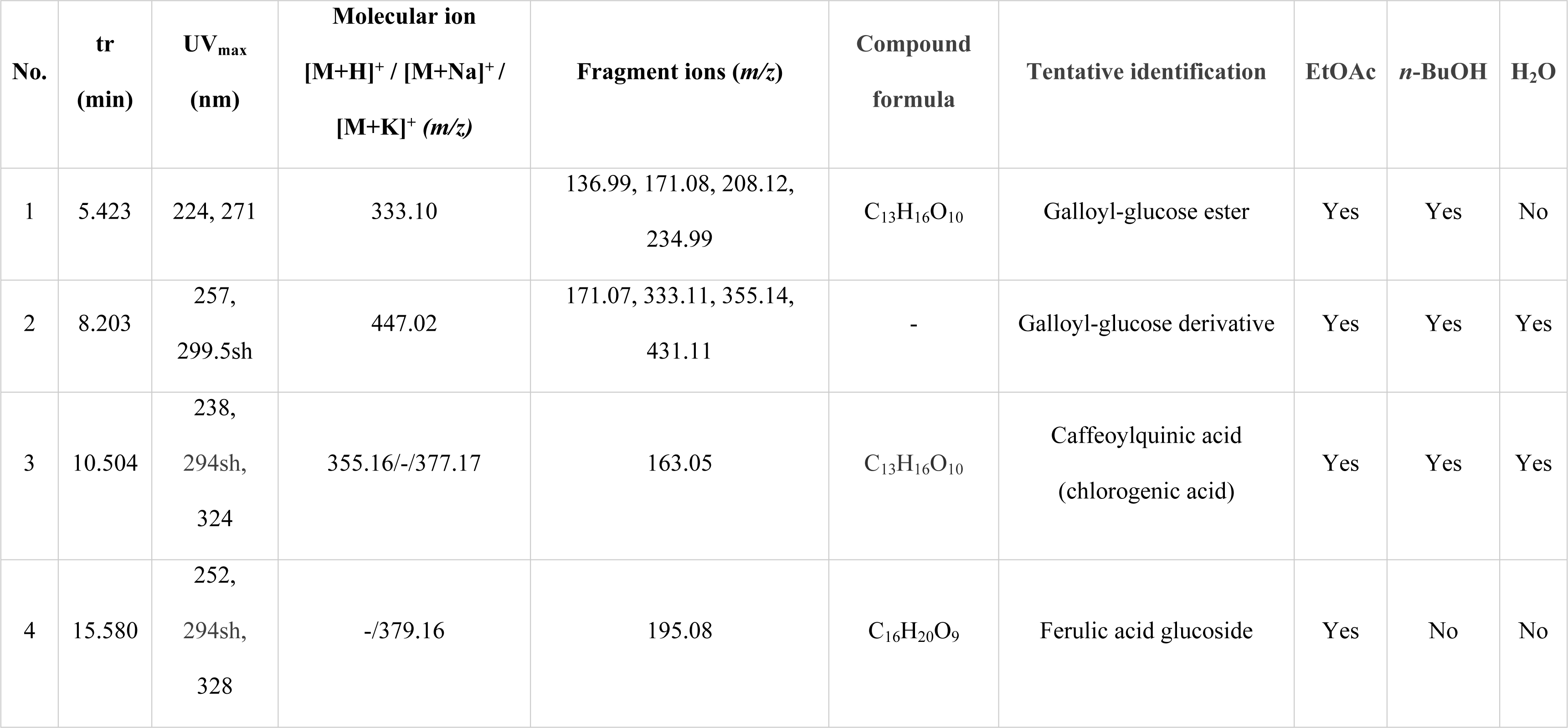
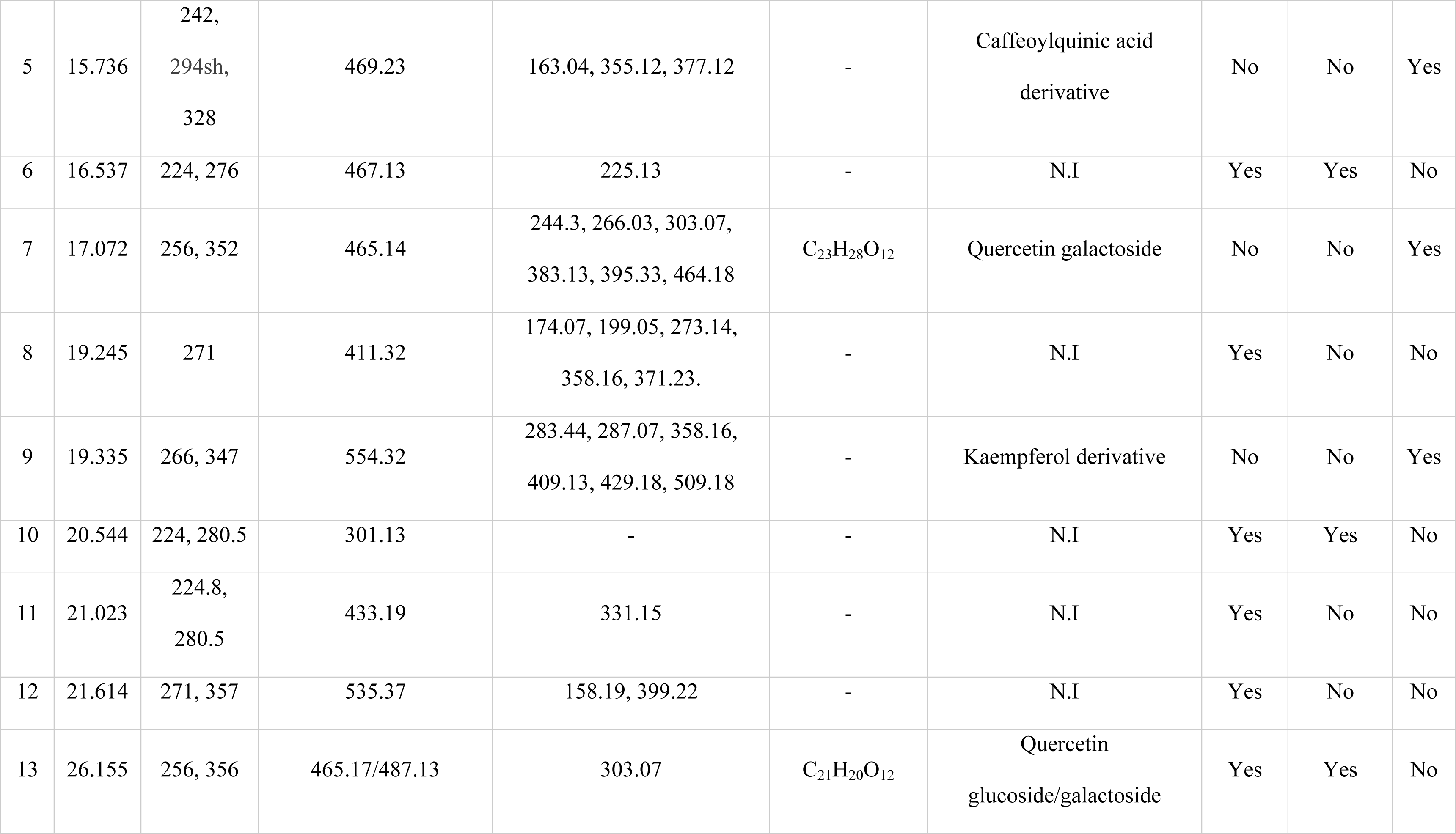
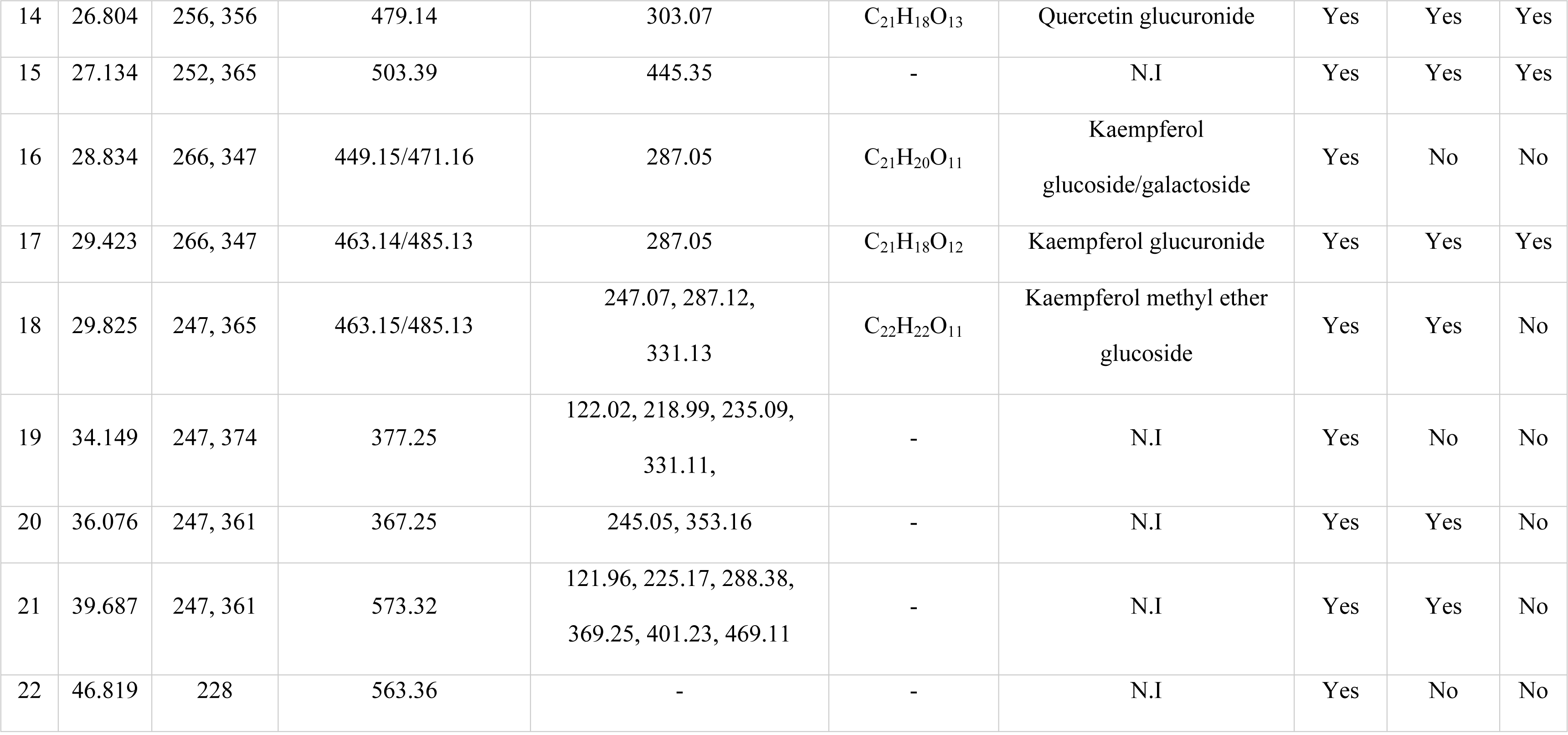
Constituents tentatively identified constituents of *E. guyoniana*.

Compounds *1* and *2* were assigned as galloyl-glucose ester and its derivative, respectively, with characteristic *m/z* 171 which corresponds to the gallic acid moiety. Compound *1* presented protonated molecular ion base peak at *m/z* 333 (C_13_H_16_O_10_H)^+^ while compound *2* presented the fragments at *m/z* 333 representing galloyl-glucose moiety and molecular ion fragment at *m/z* 447 [**28**].

Compounds *3*, *4* and *5* presented typical UV spectra of compounds with absorption bands at 220-245 nm and 315-335 nm separated by a shoulder at 290-300 nm which were characteristic of hydroxycinnamic acids [**29**].

The structure of compound *3* with [M+H]^+^ at *m/z* 355, [M+K]^+^ at *m/z* 377 and fragment at *m/z* 163 [M+H−quinic acid]^+^ corresponding to (cinnamoyl ion)^+^ was attributed to caffeoylquinic acid (chlorogenic acid) [**30**]. Compound *4* showed an UV spectrum similar to that of ferulic acid with λ_max_ at 328 nm. It also presented a pseudomolecular ion [M+Na]^+^ at m/z 379 releasing a fragment at *m/z* 195 (−162 amu, glucoside moiety) attributed to ferulic acid which allowed its tentative identification as a ferulic acid glucoside [**31, 32**]. Compound *5* with characteristic *m/z* 355 and fragment ion at *m/z* 163 (base peak) was identified as a caffeoylquinic acid derivative. The *m/z* 469 molecular ion fragment could be attributed to feruloyl caffeoyl-glycerol ([M+K]^+^) with the fragment ion m/z 377 [M+K−92]^+^ indicating the characteristic of a loss of glycerol group [**33**].

These hydrocinnamic compounds have been identified for the first time in *E. guyoniana* plant extract.

Compounds *7*, *9* and *13*-*18* presented typical absorption maxima of flavonol glycosides around 250 and 360 nm [**34, 35**].

Compounds *7*, *13* and *14* were assigned as quercetin derivatives based on their UV-DAD fingerprint (*λ*_max_ 256 and 352-356 nm) and mass spectra indicating a protonated quercetin aglycone moiety at *m/z* 303. Compounds *7* and *13* were assigned as a quercetin galactoside and quercetin glucoside/galactoside, respectively, based on their molecular ions at *m*/*z* 465 (302 + 162 (glucoside moiety) + H^+^) and 487 (302 + 162 + Na^+^) and compound *14* was identified as quercetin glucuronide with m/z 479 (302 + 176 (glucuronyl moiety) + H^+^).

All the detected quercetin derivatives were previously reported from *E. guyoniana* [**8, 11**].

Compounds *9*, *16*, *17* and *18* were assigned as kaempferol derivatives based on their mass spectra with a characteristic protonated kaempferol aglycone moiety at *m/z* 287 (286 + H) and on the UV-DAD profile (*λ*_max_ 265, and 347nm) characteristic for kaempferol. Substituents of compound *9* with molecular ion peak at *m/z* 554, detected only in the aqueous fraction, could not be determined, therefore it was identified as a kaempferol derivative. Compounds *16* and *17* exhibited [M+H]^+^ and [M+Na]^+^ at 449 (286 + 162 (glucoside moiety) + H)/471 (286 + 162 (glucoside moiety) + Na) and 463 (286 + 176 (glucuronyl moiety) + H)/485 (286 + 176 (glucuronyl moiety) + Na) were identified as kaempferol glucoside/galactoside as it was not possible to differentiate between these two and kaempferol glucuronide [**36**] respectively.

Compound *18* presented the same ion fragmentation with peaks [M+H]^+^ and [M+Na]^+^ at *m*/*z* 463 and 485, therefore it was tentatively identified as kaempferol methyl ether glucoside by comparing its UV absorbance and spectral data with previously published reports [**37**]. This compound is reported for the first time from *E. guyoniana*.

## 3. Experimental section

### 3.1 Plant material

The aerial part of *Euphorbia guyoniana* was harvested at the blooming stage, in the Wilaya of Touggourt located in the geographical coordinates: 33°06′30″ North 6°03′50″ East, Algeria, from February to March 2022. The botanical identification was performed by Dr. Lakhdari Wassima in the botanical laboratory of the High National School of Agronomy of El-Harrache (Algeria). The sample was washed in a running tap and shade-dried during 11 days until constant weight was attained. The dried sample was grounded using an industrial mill and stored in an airtight dark waterproof polyethene bag until further use.

### 3.2 Chemicals and reagents

Ethanol (96%, v/v), sodium carbonate (Na_2_CO_3_), gallic acid, quercetin, Folin–Ciocalteu reagent and aluminium chloride (AlCl_3_) were purchased from Sigma Aldrich (St. Louis, MO, USA). All the chemicals used were of analytical grade.

Methanol used for HPLC analysis (chromatographic grade) was purchased from Sigma-Aldrich (Budapest, Hungary).

### 3.3 Extraction process

A 5g aliquot of powdered sample of *Euphorbia guyoniana* was mixed with measured volumes of aqueous ethanol solvent according to the feed-to-solvent ratio variation in a 250 ml flat-bottomed flask equipped with an Allihn reflux condenser where water was used as cooling liquid. The mixture was irradiated in an enclosed reflux modified microwave oven (MW73B, Samsung, Korea: maximum delivered power of 800 Watt, frequency 2450 MHz) using 2-levels of heating: irradiation based on experimental design and 10 min of cooling. Extraction time, ethanol concentration, applied microwave power and feed-to-solvent ratio were adjusted according to the requirements of the experimental design for each run (Table 6). After the extraction process, the mixture was filtered through a Buchner glass funnel No 4 and concentrated to dryness using a rotary evaporator (Büchi Rotavapor R-200 coupled with Büchi Vac V-500 pump. Switzerland). The extracts were all dissolved in the same volume of a mixture of ethanol-water (1:1. v/v) and were stored at 4 °C in a refrigerator until further analysis.

### 3.4 Experimental design

Experiments were carried out using response surface methodology (RSM) to obtain optimized extraction conditions for dependent variables: total phenolic content and total flavonoid content. A central composite design (CCD) was employed with four independent variables in five different coded levels (-α, -1, 0, +1, +α). The CCD was selected as it provides, besides of the center points, a group of “star points” that represent lower and higher extreme values of the design and thus allow estimation of curvature and augmentation of the variable domain from -α to +α [**38**]. The α values were of -2 and +2 and were calculated according to the following Eq. (3):

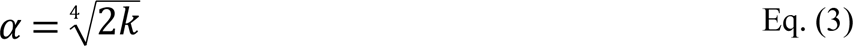

where k is the number of independent variables

The four independent variables, i.e. extraction time (*x_1_*), ethanol concentration in the extractive solvent (*x_2_*), microwave power (*x_3_*) and feed-to-solvent ratio (*x_4_*) in the five coded levels are presented in Table 5.

**Table 5.** Input variables and their respective levels used to investigate the efficiency of microwave-assisted extraction.

| Independent variables | Codes | Domain | Levels |  |  |  |  |
| --- | --- | --- | --- | --- | --- | --- | --- |
| | | | $-\alpha$ | -1 | 0 | +1 | $+\alpha$ |
| **Extraction time (min)** | $x_1$ | 5 – 25 | 5 | 10 | 15 | 20 | 25 |
| **Ethanol concentration (%)** | $x_2$ | 30 – 70 | 30 | 40 | 50 | 60 | 70 |
| **Microwave power (Watt)** | $x_3$ | 180 – 800 | 180 | 300 | 450 | 600 | 800 |
| **Feed-to-solvent ratio (g/ml)** | $x_4$ | 1:7.5 – 1:17.5 | 1:7.5 | 1:10 | 1:12.5 | 1:15 | 1:17.5 |

A second order polynomial equation was postulated for each response Y following Eq. (4):

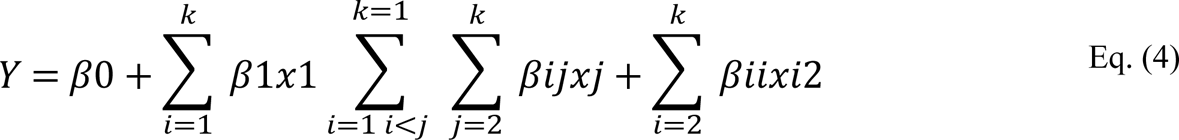

where *x_i_*values are independent parameters affecting the dependent responses Y

β0, βi, βii, and βij are the regression coefficients for intercept, linear, quadratic and interaction terms respectively

Table 6 indicates the CCD experimental data which consists of 27 combinations including three replicates at the central point with the four variables and the two responses.

**Table 6.**
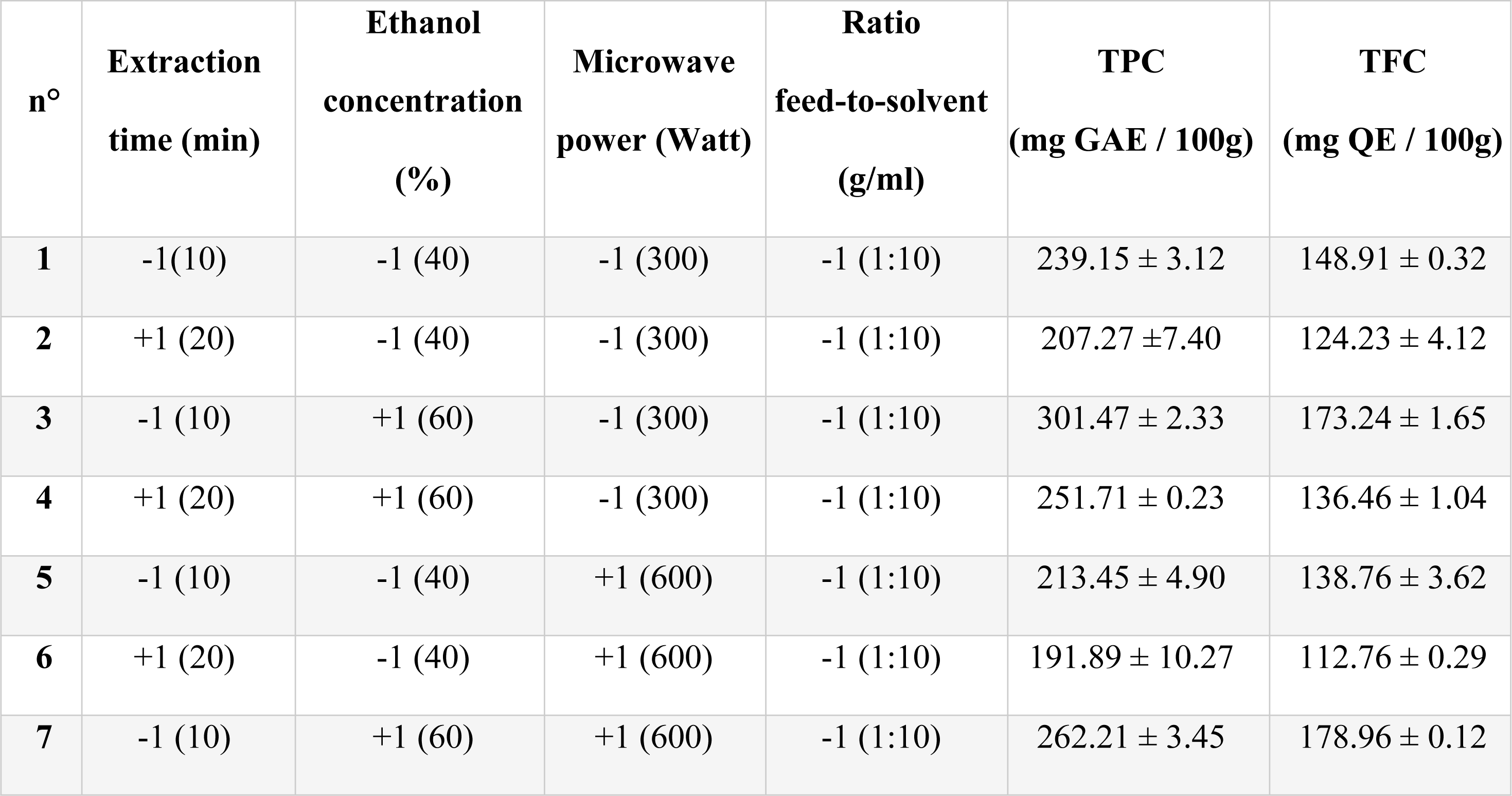
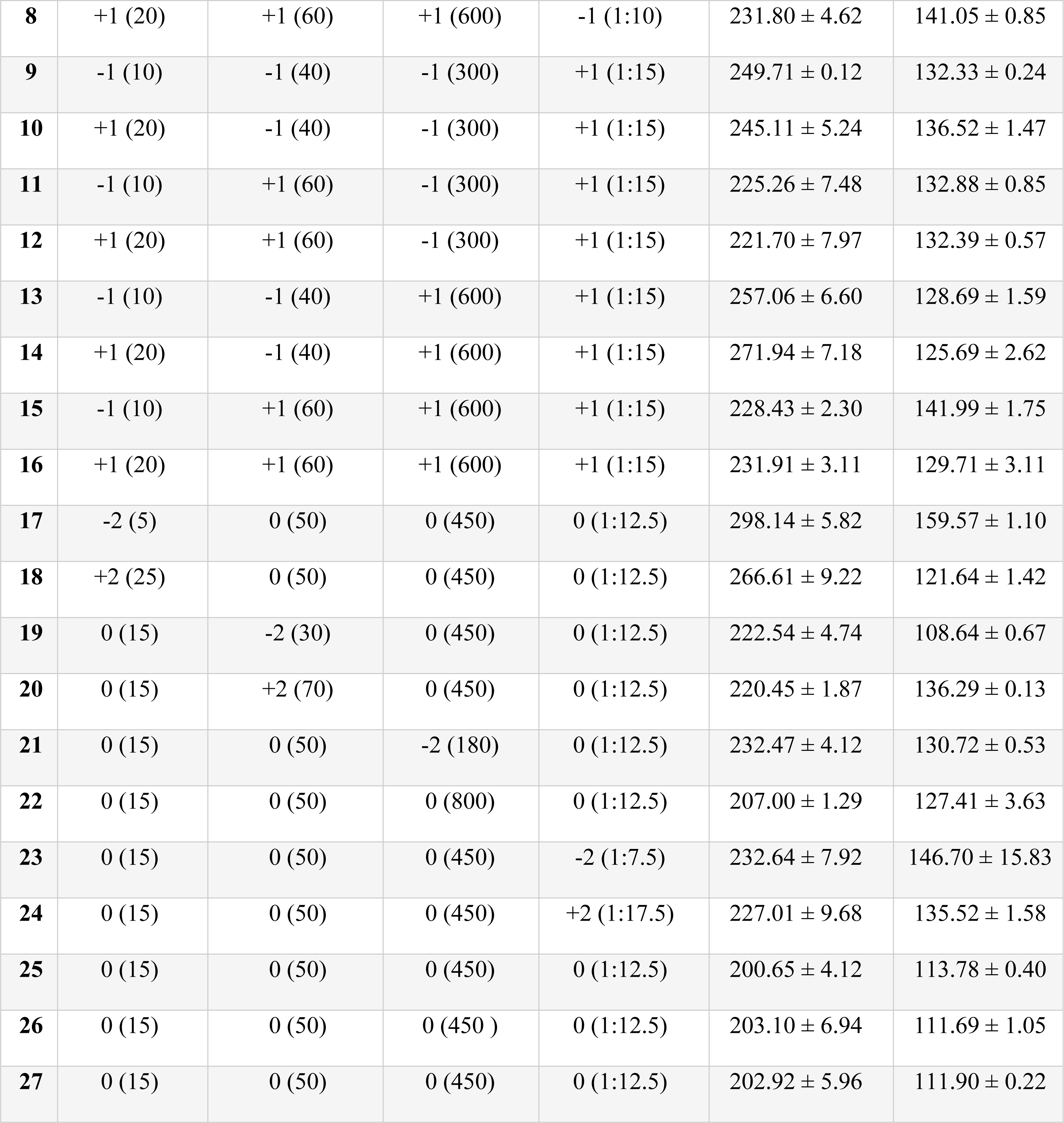
Central composite design and calculated results.

### 3.5 Total Phenolic Content (TPC) Determination

The Total Phenolic Content in each *Euphorbia guyoniana* extract was determined using the Folin–Ciocalteu’s colorimetric method with slight modification [**39, 40**]. Each extract was diluted 50 times prior to the analysis. An aliquot of 0.5 ml of each diluted extract was mixed with 2 ml of Folin–Ciocalteu reagent at 2M, diluted 10 times in distilled water. Finally, 2.5 ml of sodium carbonate at 2% was added. The mixture was heated at 45°C during 15 min. The absorbance was spectrophotometrically measured at 765 nm against blank. The obtained results were expressed as milligram gallic acid equivalent per 100 grams (mg GAE/100g) based on the standard calibration curve of gallic acid (0 − 180 mg/l, y = 135.12x + 10.238, R^2^ = 0.993). All assays were carried out in triplicates and the results expressed as mean values ± standard error.

### 3.6 Total Flavonoid Content (TFC) Determination

The Total Flavonoid Content in each *Euphorbia guyoniana* extract was determined using aluminum chloride (AlCl_3_) colorimetric method with slight modification [**41**]. Each extract was diluted 100 times prior to the analysis. An aliquot of 1 ml of each diluted extract was mixed with 2 ml of AlCl_3_ at 2%. The mixture was kept at room temperature for 15 min and the absorbance was measured at 420 nm against blank. The obtained results were expressed as milligram quercetin equivalent per 100 grams (mg QE/100g) based on the standard calibration curve of quercetin (0 – 40 mg/l, y = 32.155x + 0.1281, R^2^ = 0.9994). All assays were carried out in triplicates and the results expressed as mean values ± standard error.

All absorbance readings were performed on UV-vis Spectrophotometer (UV mini 1240, Shimadzu, Japan).

### 3.7 *In vivo* anti-inflammatory activity: Carrageenan rat paw edema model

The *in vivo* anti-inflammatory activity of the optimal extract of *Euphorbia guyoniana* was determined using the rat paw-edema assay [**42**] using a dosage of 50 mg/kg.

Ethical approval for the animal studies was obtained from the Ethics Committee for Animal Experimentation of the Faculty of Natural and Life Sciences, Department of Biology, Saad Dahleb University, Blida 1, Blida, Algeria, reference number: FSNV012023. Throughout our study, ethical standards in animal experimentation were adhered to in accordance with the EU Directive 2010/63/EU. All applicable international, national, and/or institutional guidelines for the care and use of animals were followed.

In the present study, 18 Wistar rats weighing between 220 and 250 g were purchased from the Pasteur Institute of Algiers (IPA, Algiers, Algeria). The animals were maintained under standard conditions with access to distilled water and pelleted food ad libitum during two weeks (adaptation period). Prior to the anti-inflammatory study, the animals were kept under fasting conditions (no food only) for 18 hours. The animals were divided into 3 groups of 6 rats each. The control group was treated with physiological saline (Group 1). The group 2 was treated with ibuprofen (50 mg/kg) as the reference drug (positive control group). The group 3 was orally administered with 50 mg/kg dose of optimal extract of *Euphorbia guyoniana*.

After 30 min, 0.1 ml of 0.5% carrageenan suspension in saline was injected subcutaneously into the plantar aponeurosis of the left hind paw. The monitoring of edema volume was using plethysmometer for 4 hours. Paw volume was measured before the carrageenan injection, that is, at “0 hour” and then at 1, 2, 3 and 4 hour after carrageenan injection.

The percentage increase in paw edema and the percentage of edema inhibition were calculated as followed in the Eqs. (5) and (6):

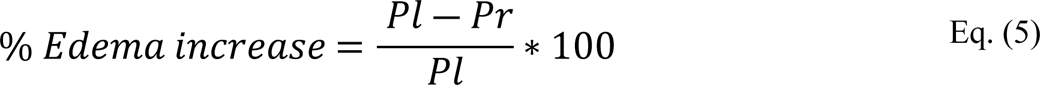

where P_l_: means weight of the left paws;

P_r_: means weight of the right paws

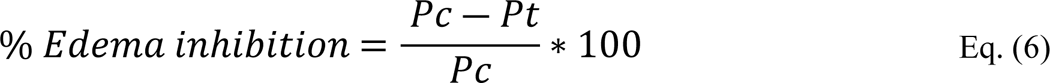

where P_c_: average edema weight of the negative control group;

P_t_: average edema weight of the tested group

### 3.8 Phenolic compounds profile via RP-HPLC-PDA-ESI-MS

Phenolic compounds profile was realized by using a Waters 2695 (Milford, MA) HPLC system equipped with Waters 2988 Photodiode Array and coupled to a Waters ACQUITY QDA with an electrospray ionization source (ESI). Separation of phenolic compounds was carried out on Phenomenex (Torrance, CA, USA) Kinetex 5μm XB-C18A (100 Å 250 x 4.6mm) at 30°C, using a binary mobile phase composed of acidified water (1% v/v acetic acid; A) and methanol (B). The following gradient program was applied at a flow rate of 1 ml/min: 0–1 min 95% A; 1–55 min 5-95% B; 55–57 min 5-95% A; 57–65 min 95% A with linear gradient and UV-detection from 210 to 400 nm. MS analysis was carried out using positive ion mode at voltage of 0.8 Kv in full Scan mode (*m/z* 50–600).

### 3.9 Statistical analysis

The complete response surface methodology experimental design and the analysis of variance using Fisher test (ANOVA, F-test) were generated by a statistical software package Design Expert 13 software^®^ (Version 13.0.5.0. Stat-Ease Inc. Minneapolis, USA). The ANOVA was used to analyze and identify the significant factors, and subsequently, the accuracy of the suggested models was determined. Also, p-value (p ≤ 0.05), the “lack of fit”, the coefficient of determination R^2^, the adjusted R^2^ and the predicted R^2^, the adequate precision and the coefficient of variation (C.V.) were used to estimate the quality of the fit of the polynomial models. The desirability function approach was applied for the optimization of the extraction conditions. The validation of obtained models were performed by comparing the predicted and experimental contents under the optimized conditions. Experimental results under applied optimal conditions were performed in triplicate and the values obtained were presented as mean ± standard deviation. The 3D plot of the parameters were used to establish the model equation.

## 4. Conclusion

Response surface methodology using central composite design was successfully applied to study and optimize the influence of microwave-assisted extraction conditions on the total phenolic content and total flavonoid content of *Euphorbia guyoniana* extract. The optimal extraction conditions were found to be irradiation time of 25 min, microwave power of 450 Watt, concentration of ethanol in the extractive solvent of 40.57% and feed-to-solvent ratio of 1:17.5 g/mL resulting in total phenolic content estimated to be 376.16 mg GAE/100g and total flavonoid content to be 188.46 mg QE/100g. These experimental values were in good agreement with the predicted values, therefore it may be concluded that the generated mathematical models adequately reflected the behavior of the selected parameters for the extraction of bioactive compounds from *E. guyoniana*.

The *in vivo* anti-inflammatory activity of the optimal extract of the plant was determined using the carrageenan-induced rat paw-edema assay. A higher percentage of inhibition of paw edema of 75.98% was observed when compared with a reference treatment, i.e., ibuprofen (positive control).

The extract was analyzed by RP-HPLC-PDA-ESI-MS that revealed the abundant presence of hydroxycinnamic acids and flavonol glycosides. Among the identified phenolic compounds, galloyl-glucose, galloyl-glucose derivative, caffeoylquinic acid and its related derivative, ferulic acid glucoside and kaempferol methyl ether glucoside are reported for the first time in *E. guyoniana* extract.

To the best of our knowledge, this is the first report on the microwave-assisted extraction and optimization of *Euphorbia guyoniana* and the first evidence on the anti-inflammatory effect of *E. Guyoniana*, which supports its traditional Moroccan and Algerian use as a local treatment of venomous bites and stings. Our results therefore pave the way towards the development of innovative, evidence-based therapeutic applications of a long-used ethnomedicine, *Euphorbia guyoniana* in the management of skin disorders as it is already the case for many different Euphorbia species.

## Acknowledgements

The authors would like to thank Dr. Wassima Lakhdari from the botanical laboratory of the High National School of Agronomy of El-Harrache (Algeria) for providing the starting raw plant material and for her expertise in the botanical identification and the Laboratory of Biology and Physiology of Organisms of the University of Sciences and Technology Houari Boumediene for technical support

## Supporting information

**S1 Table. Experimental and Predicted values of total phenolic content and total flavonoid content**

**S1 Fig. Chromatogram of ethanolic extract of aerial plant extract of *Euphorbia Guyoniana***

**S2 Fig. Chromatogram of aqueous extract of aerial plant extract of *Euphorbia Guyoniana***

**S3 Fig. Chromatogram of buthanolic extract of aerial plant extract of *Euphorbia Guyoniana***

